# α-Synuclein pre-formed fibrils induce prion-like protein aggregation and neurotoxicity in *C. elegans* models of Parkinson’s disease

**DOI:** 10.1101/2021.11.11.468276

**Authors:** Merry Chen, Julie Vincent, Alexis Ezeanii, Saurabh Wakade, Shobha Yerigenahally, Danielle E. Mor

## Abstract

Parkinson’s disease (PD) is a debilitating neurodegenerative disorder characterized by progressive motor decline and the aggregation of α-synuclein protein. Growing evidence suggests that α-synuclein aggregates may spread from neurons of the digestive tract to the brain in a prion-like manner. While rodent models have recapitulated gut-to-brain α-synuclein transmission, animal models that are amenable to high-throughput investigations are needed to facilitate the discovery of disease mechanisms. Here we describe the first *C. elegans* models in which feeding with α-synuclein pre-formed fibrils (PFFs) induced prion-like dopamine neuron degeneration and seeding of aggregation of human α-synuclein expressed in the host. PFF acceleration of α-synuclein aggregation in *C. elegans* muscle cells was associated with a progressive motor deficit, whereas feeding with α-synuclein monomer produced much milder effects. RNAi-mediated knockdown of the *C. elegans* syndecan *sdn-1*, and enzymes involved in heparan sulfate proteoglycan biosynthesis, afforded protection from PFF-induced seeding of aggregation and toxicity, as well as dopaminergic neurodegeneration. This work offers new models by which to investigate gut-derived α-synuclein spreading and propagation of disease.

## Introduction

Parkinson’s disease (PD) is a debilitating movement disorder characterized by loss of dopaminergic neurons and formation of Lewy body inclusions containing aggregated α-synuclein protein^1-2^. While the etiology of the disease remains unknown, growing evidence suggests that α-synuclein aggregation may originate in neurons that innervate the gastrointestinal tract, and spread to the central nervous system (CNS) via a prion-like mechanism^3-4^. Postmortem studies have found α-synuclein pathology in the enteric nervous system during early stages of PD^5^, and gastrointestinal dysfunction is often among the first disease symptoms^6^. Rodent models have recapitulated the spread of α-synuclein pathology with associated neurodegeneration and motor deficits following oral administration^7^ or gastrointestinal injection^8-9^ of recombinant α-synuclein pre-formed fibrils (PFFs). However, it remains unclear how α-synuclein originating in the digestive tract spreads to the central nervous system and induces neurodegeneration.

*C. elegans* offers a unique model system by which to address these questions, having orthologs for 60-80% of human genes, exceptionally high genetic tractability, and suitability to rapid, large-scale behavioral and phenotypic screening approaches^10-11^. The worm nervous system uses highly conserved signaling components to give rise to a complex set of behaviors^10^. Similar to humans, *C. elegans* has an alimentary nervous system that facilitates feeding and acts largely independently from neurons in the rest of the body (termed the somatic nervous system)^12^. While some studies using *C. elegans* have shown α-synuclein cell-to-cell transmission within the somatic nervous system^13^, or between neurons and other tissues^14-15^, it remains to be demonstrated whether α-synuclein of digestive origin can seed the aggregation of α-synuclein localized in body tissues and promote neurodegeneration in a prion-like manner.

Here, we have generated the first such models in *C. elegans*, by feeding worms human α-synuclein PFFs. We show that in animals expressing human α-synuclein (‘host α-synuclein’), PFF feeding accelerates dopamine neuron degeneration and promotes α-synuclein aggregation in muscle cells. We find that several members of the heparan sulfate proteoglycan biosynthesis pathway mediate both PFF-induced α-synuclein aggregation and neurodegeneration, providing a novel platform by which to investigate prion-like activity of α-synuclein and neurotoxic mechanisms.

## RESULTS

### Prion-like spreading of α-synuclein from the digestive tract to neurons causes dopaminergic neurodegeneration

Prions and prion-like proteins, including α-synuclein, exhibit self-templated replication or ‘seeding’ via the repeated conversion of soluble protein into misfolded/ aggregated conformations. This process requires endogenous protein as a source of aggregate amplification^16^. Since *C. elegans* does not naturally possess an α-synuclein homolog, we took advantage of existing transgenic strains expressing human wild-type α-synuclein that exhibit mild degenerative phenotypes^17-18^, and asked whether α-synuclein PFF feeding could accelerate disease.

Recombinant α-synuclein PFFs were prepared using standard methods^19^, and fibrillization was monitored by Thioflavin T and sedimentation analysis to confirm that the protein had become insoluble (Fig. S1). To facilitate uptake of PFFs by ingestion, we pre-mixed sonicated PFFs with the bacterial food source, and treated animals on day 1 of adulthood. Adult-only treatment was also chosen to avoid potential developmental effects of PFF toxicity.

Following only 24 hours of incubation with PFFs from day 1 to day 2, worms expressing pan-neuronal human wild-type α-synuclein^17^ exhibited a significant loss of CEP dopamine neurons compared with buffer-treated animals, as measured on day 5 of adulthood (Fig. 1A, B). ADE dopamine cell bodies were spared (Fig. 1A, C). Prion-like seeding of aggregation requires that the initial seed be an aggregated or misfolded conformer, rather than a physiological monomer^16^. Therefore, we tested α-synuclein monomer treatment, and found that this was insufficient to cause CEP cell body loss (Fig. 1A, B). Similarly, PFF treatment of worms not expressing α-synuclein had no effect on dopamine neurons, as expected (Fig. 1A-D). We did observe that dopaminergic neurites were vulnerable to both PFF and monomer treatments (Fig. 1A, D), suggesting that neurites may be particularly sensitive to increased concentration of α-synuclein, irrespective of its aggregation state.

**Fig. 1:**
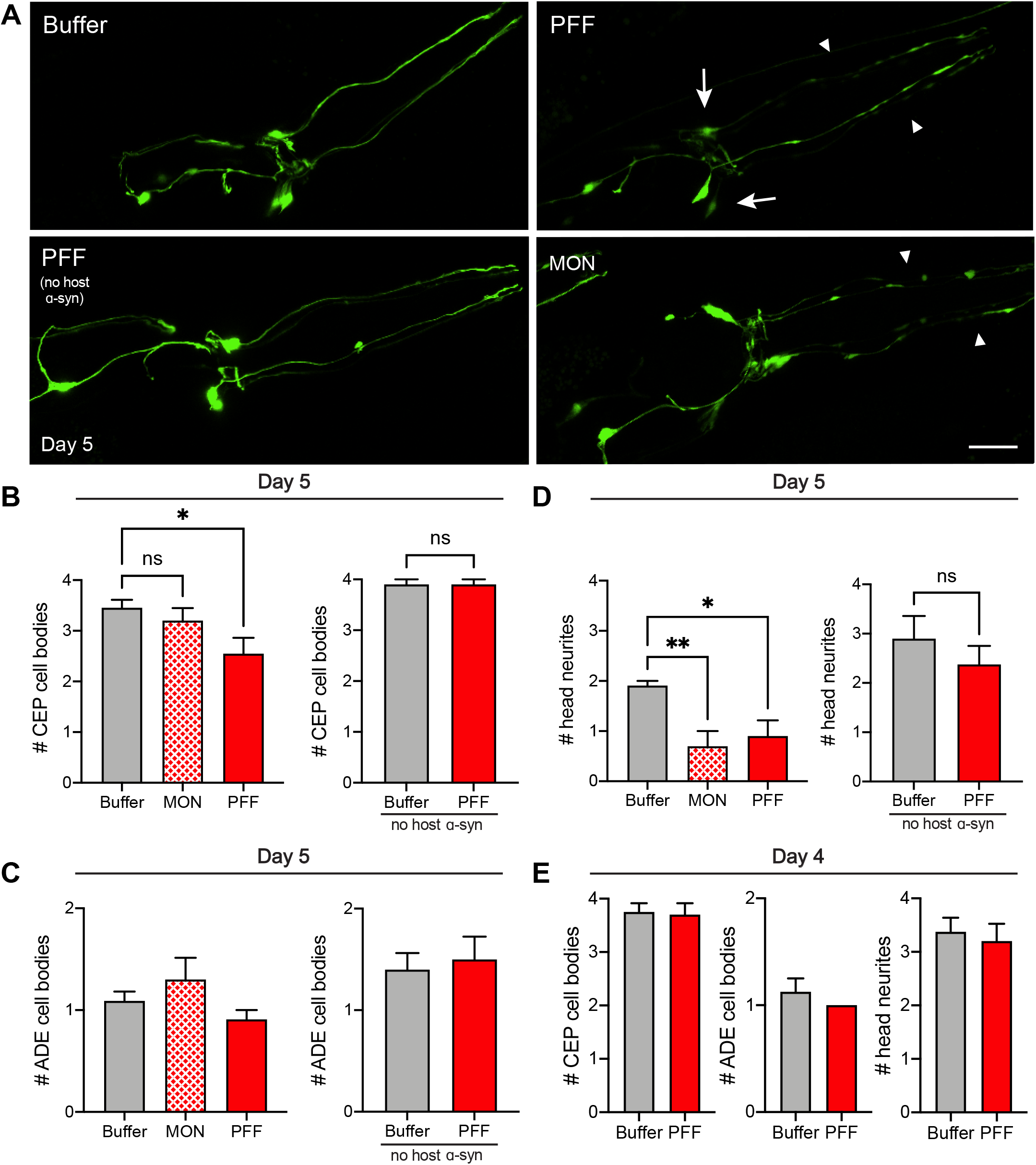
α-synuclein PFF feeding accelerates dopaminergic neurodegeneration. **A**, Worms expressing pan-neuronal human wild-type α-synuclein were treated with 0.25 ng/μL PFFs or monomer (MON), or buffer alone, on day 1 of adulthood. Worms expressing GFP in dopamine neurons without transgenic α-synuclein expression were also fed PFFs on day 1. On day 5, dopamine neurons were imaged and scored for degenerating cell bodies (arrows) and neurites (arrowheads). Scale bar, 20 μm. **B**, CEP cell body quantification in worms expressing neuronal α-synuclein (left panel), or not (right panel). Data are mean±s.e.m. **C**, ADE cell body quantification in worms expressing neuronal α-synuclein (left panel), or not (right panel). Data are mean±s.e.m. **D**, Neurite quantification in worms expressing neuronal α-synuclein (left panel), or not (right panel). Data are mean±s.e.m. In B-D, n=11 each for Buffer, PFF in left panel, n=10 for each of the other groups. Left panels, One-Way ANOVA with Tukey’s post-hoc. Right panels, Two-tailed *t*-test. **E**, Dopamine neuron scoring on day 4 in worms expressing neuronal α-synuclein. Data are mean±s.e.m. n=8 each for Buffer, 10 each for PFF. Two-tailed *t*-tests. ns, not significant. \**P*<0.05, \*\**P*<0.01.

To determine if the loss of CEP cell bodies due to PFF feeding is age-dependent, we quantified the extent of dopamine neuron degeneration one day earlier, on day 4. Unlike our observations on day 5, worms expressing neuronal α-synuclein did not show additional damage to CEP cell bodies or neurites with PFF feeding (Fig. 1E). These results indicate that gut-derived PFFs induce dopamine neurodegeneration in an age-dependent manner, mimicking the late-onset development of PD and consistent with the long incubation times of prion diseases in humans^16^.

### Gut-derived α-synuclein pre-formed fibrils promote toxic aggregation of host α-synuclein

To test if PFFs of digestive origin are able to promote aggregation of α-synuclein in body tissues, we used a well-established model in which human wild-type α-synuclein fused to YFP is expressed in muscle cells for easy visualization of aggregates^18^. PFF feeding in this strain resulted in a significant increase in the number of aggregates (Fig. 2A, B) as well as the total aggregate area (Fig. 2A, C) as early as day 2 of adulthood. Consistent with prion-like mechanisms, monomeric α-synuclein did not increase aggregate load (Fig. 2A-C), and expression of YFP alone in muscle cells (no host α-synuclein) showed no aggregate-like puncta (Fig. 2A-C). These findings suggest that PFFs are able to rapidly spread from the digestive tract to body tissues, and seed the aggregation of host α-synuclein at these distant sites.

**Fig. 2:**
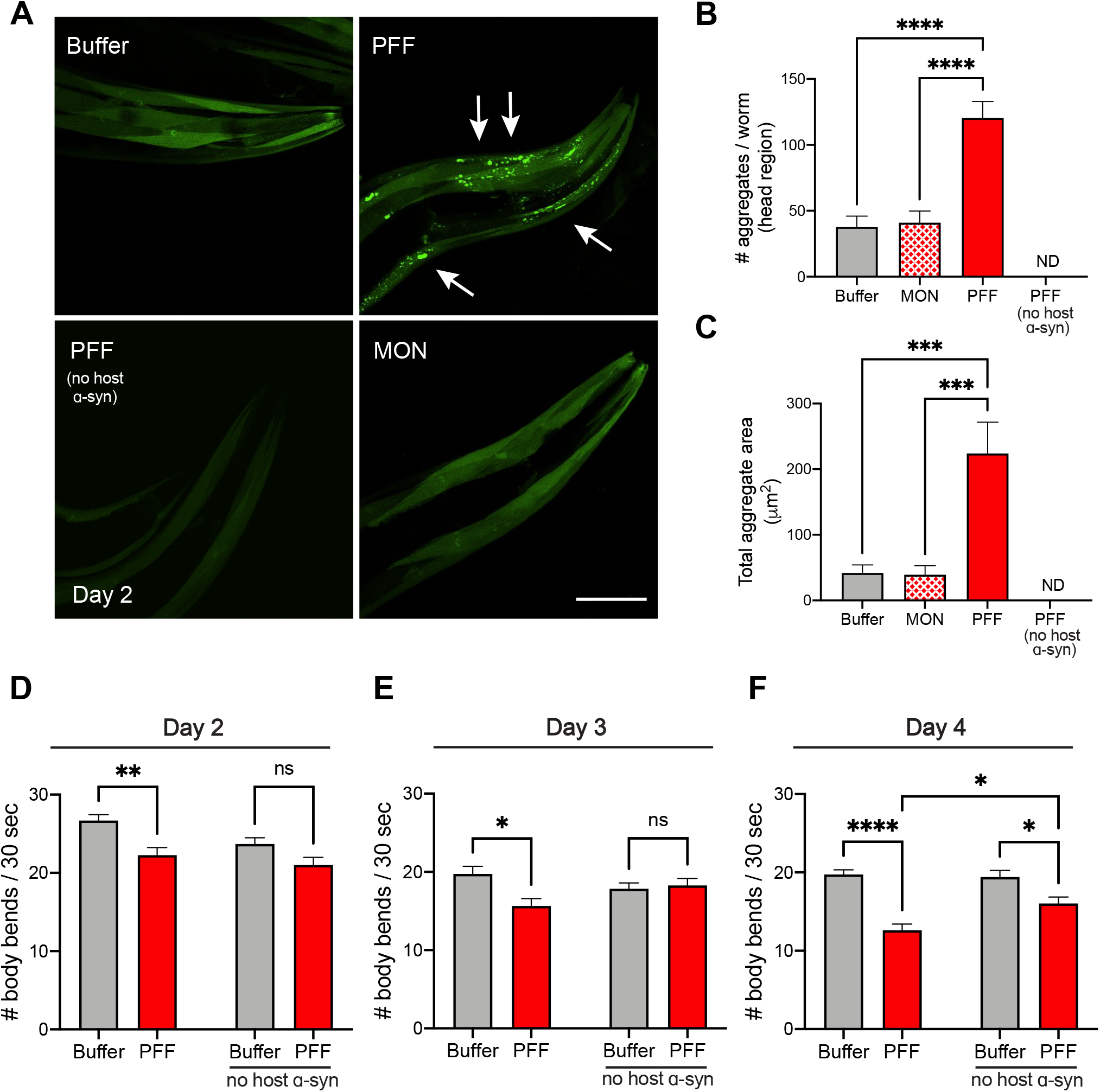
PFF feeding promotes aggregation of host α-synuclein and associated motor decline. **A**, Worms expressing human wild-type α-synuclein-YFP in muscle cells were fed 0.25 ng/μL PFFs or monomer (MON), or buffer alone, on day 1 of adulthood. Worms expressing YFP in muscle without transgenic α-synuclein expression were also fed PFFs on day 1. On day 2, α-synuclein aggregates (arrows) were imaged. Scale bar, 50 μm. **B**, Quantification of # α-synuclein aggregates per worm. ND, not detected. Data are mean±s.e.m. One-Way ANOVA with Tukey’s post-hoc. **C**, Quantification of total aggregate area. ND, not detected. Data are mean±s.e.m. One-Way ANOVA with Tukey’s post-hoc. In B-C, n=22 for Buffer, 23 for PFF, 21 for MON, and 11 for PFF (no host α-synuclein). **C**, Motor assays on day 2 for α-synuclein-YFP and non-transgenic strains. Data are mean±s.e.m. n=19 for each group. Two-Way ANOVA with Sidak’s post-hoc. **D**, Motor assays on day 3 for α-synuclein-YFP and non-transgenic strains. Data are mean±s.e.m. n=19 for each group. Two-Way ANOVA with Sidak’s post-hoc. **E**, Motor assays on day 4 for α-synuclein-YFP and non-transgenic strains. Data are mean±s.e.m. n=38 for each group. Two-Way ANOVA with Sidak’s post-hoc. ns, not significant. \**P*<0.05, \*\**P*<0.01, \*\*\**P*<0.001, \*\*\*\**P*< 0.0001.

We next asked whether increased aggregation is associated with toxicity. To address this, we conducted crawling assays as a measure of basic motor function. As early as day 2, and concomitant with increased aggregation (Fig. 2A-C), we detected a significant reduction in motility of PFF-fed worms expressing α-synuclein in muscle, compared with buffer-treated animals (Fig. 2D). In contrast, non-transgenic worms did not show a deficit at this time-point despite PFF feeding (Fig. 2D), suggesting the toxicity is dependent on host α-synuclein.

To test if the motor impairment is progressive, we examined days 3 and 4, and found that worms expressing muscle α-synuclein continue to have a decline in motor function that is most pronounced on day 4 (Fig. 2E-F). While non-transgenic worms fed PFFs do not show motor dysfunction on day 3, by day 4 these animals do exhibit a reduction in motility (Fig. 2E-F). The non-prion-like mechanisms of PFF toxicity occurring in non-transgenic worms by this time-point could not account, however, for the entire loss of motor function in animals that do express host α-synuclein. Indeed, we detected a significant difference between PFF-treated animals with and without host α-synuclein expression (Fig. 2F), indicating that on day 4 a combination of both prion- and non-prion-like mechanisms contribute to PFF-induced behavioral decline.

To further test whether PFF toxicity is dose-dependent, PFF concentrations from 0.1 ng/μL to 1.0 ng/μL were fed to worms expressing α-synuclein in muscle, and crawling behavior was measured on days 2, 3, and 4 (Fig. 3). Whereas the lowest dose, 0.1 ng/μL was insufficient to cause a loss of motor function (Fig. 3A), 0.25 ng/μL PFF induced a steady decline in movement that was significantly different from buffer-treated controls (Fig. 3B). This dosage matches that of our neurodegeneration and aggregation experiments (Fig. 1-2), and consistently showed progressive deterioration of motor function over time. Treatment with 0.25 ng/μL monomer was not significantly different from control, but was significantly less toxic than the same dosage of PFFs (Fig. 3B), consistent with prion-like underlying mechanisms. Interestingly, higher concentrations of PFFs (0.5 and 1.0 ng/μL) each induced a severe decline in motor function immediately by day 2, and did not show progressive worsening thereafter (Fig. 3C-D). Matching monomer concentrations acted similarly to PFFs at these high dosages, causing significantly worse functioning than that observed in buffer-treated controls (Fig. 3C-D). These findings suggest that PFF dosages of 0.5 ng/μL or higher cause severe outcomes that rely on fundamentally different mechanisms than the 0.25 ng/μL dose, which is more relevant to the study of prion-like α-synuclein toxicity.

**Fig. 3:**
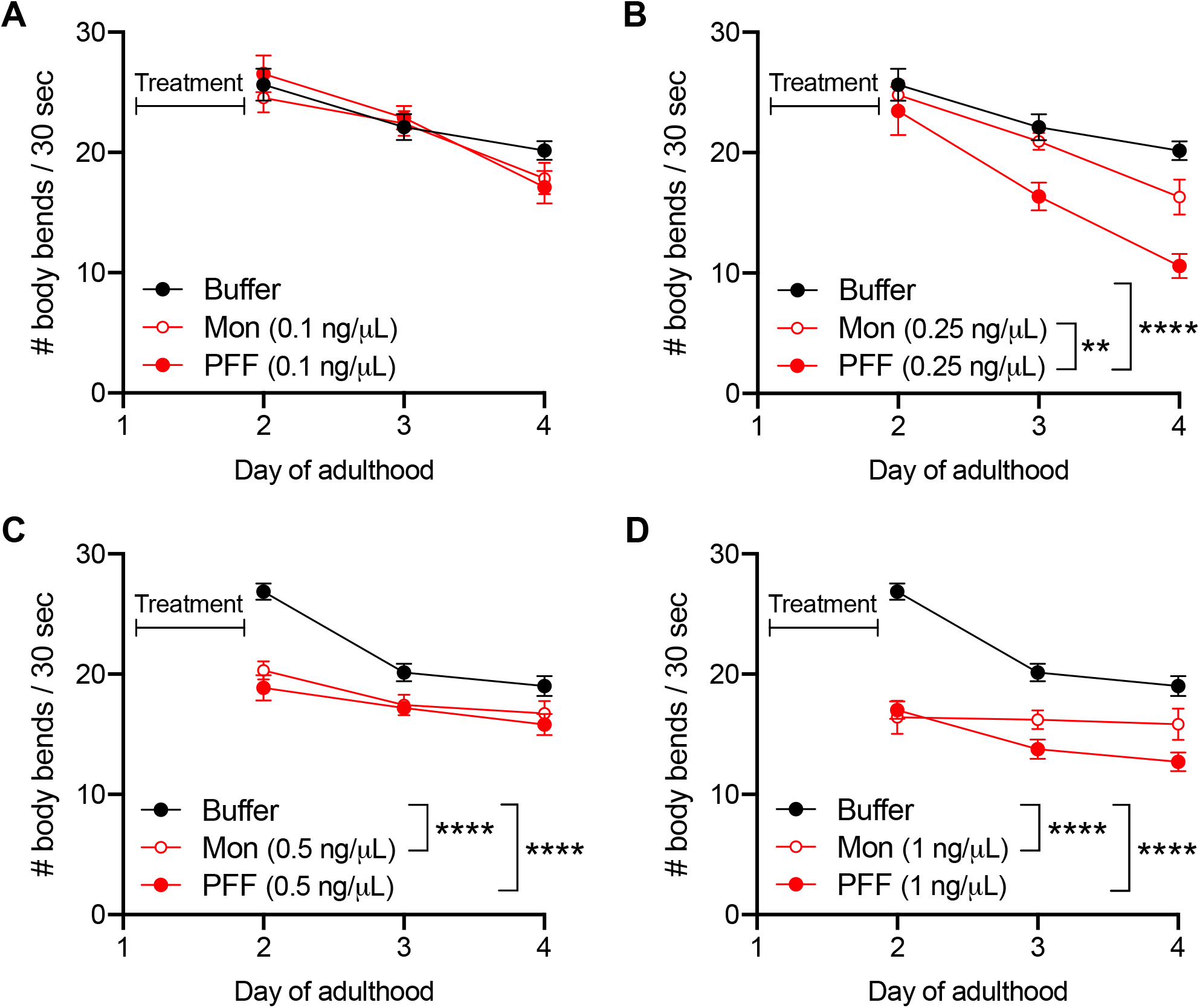
Dose-dependent toxicity of PFF feeding. **A**, Worms expressing human wild-type α-synuclein-YFP in muscle cells were fed 0.1 ng/μL PFFs or monomer (MON), or buffer alone, on day 1 of adulthood and motor assays were conducted on days 2, 3, and 4. Data are mean±s.e.m. n=19 for each group. Two-Way ANOVA with Tukey’s post-hoc. **B**, Worms expressing human wild-type α-synuclein-YFP in muscle cells were fed 0.25 ng/μL PFFs or monomer (MON), or buffer alone, on day 1 of adulthood and motor assays were conducted on days 2, 3, and 4. Data are mean±s.e.m. n=19 for each group. Two-Way ANOVA with Tukey’s post-hoc. **C**, Worms expressing human wild-type α-synuclein-YFP in muscle cells were fed 0.5 ng/μL PFFs or monomer (MON), or buffer alone, on day 1 of adulthood and motor assays were conducted on days 2, 3, and 4. Data are mean±s.e.m. n=38 for each group. Two-Way ANOVA with Tukey’s post-hoc. **D**, Worms expressing human wild-type α-synuclein-YFP in muscle cells were fed 1.0 ng/μL PFFs or monomer (MON), or buffer alone, on day 1 of adulthood and motor assays were conducted on days 2, 3, and 4. Data are mean±s.e.m. n=38 for each group. Two-Way ANOVA with Tukey’s post-hoc. \*\**P*<0.01, \*\*\*\**P*< 0.0001.

### α-Synuclein prion-like transmission from the digestive tract to other tissues is mediated by heparan sulfate proteoglycans

To uncover potential regulators of α-synuclein neurotoxic spread, we performed a targeted screen of eight heparan sulfate proteoglycan (HSPG) pathway genes. While the precise mechanisms by which gut-derived α-synuclein PFFs enter other tissues is not known, evidence from mammalian cell culture studies suggests that PFFs can interact with HSPG cell surface receptors and become endocytosed^20^. To test if HSPGs may play a role in our new PFF-based *C. elegans* models of PD, worms expressing human α-synuclein in muscle were fed RNAi corresponding to each HSPG gene starting at the L4 stage, such that gene knockdown would occur after development and by the time animals were fed 0.25 ng/μL PFFs on day 1 of adulthood. On day 4, the time-point at which we had documented the greatest reduction in movement behavior of PFF-fed animals at this dosage (Fig. 2F, 3B), motor function was tested using the crawling assay. As expected, PFF treatment on empty vector RNAi significantly reduced motility compared with buffer-treated vector RNAi controls (Fig. 4A).

**Fig. 4:**
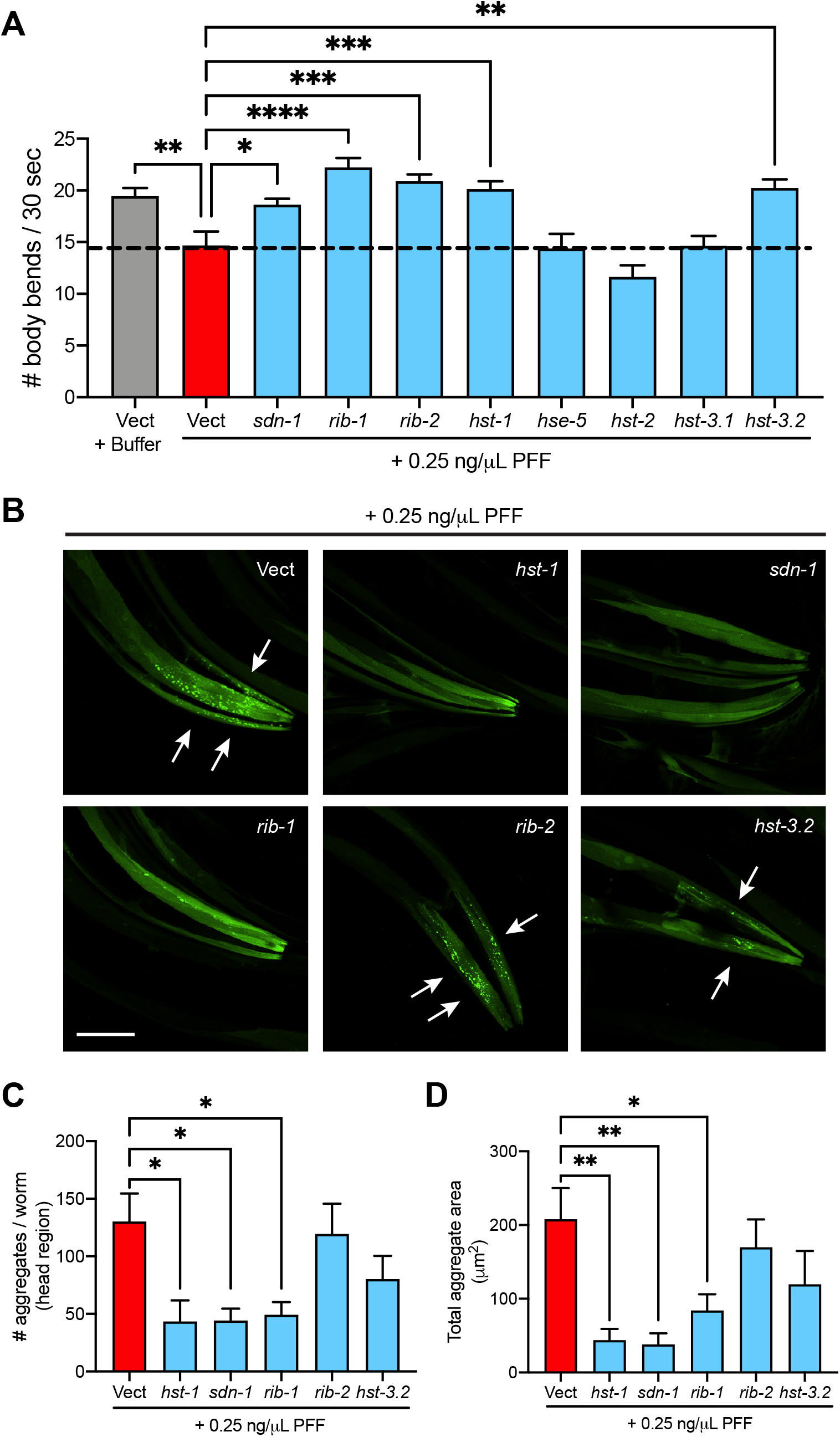
HSPG gene knockdown rescues motor function and reduces α-synuclein aggregation in PFF-fed animals. **A**, Worms expressing human wild-type α-synuclein-YFP in muscle cells were fed 0.25 ng/μL PFFs or buffer on day 1 of adulthood and motor assays were conducted on day 4. Data are mean±s.e.m. n=38 for *hst-1* and 19 for each of the other groups. One-Way ANOVA with Dunnett’s post-hoc. **B**, On day 4, α-synuclein aggregates (arrows) were imaged. Scale bar, 50 μm. **C**, Quantification of # α-synuclein aggregates per worm. Data are mean±s.e.m. One-Way ANOVA with Dunnett’s post-hoc. **D**, Quantification of total aggregate area. Data are mean±s.e.m. One-Way ANOVA with Dunnett’s post-hoc. In C-D, n=15 for Vect, 13 for *rib-1*, 10 for *hst-3*.*2*, and 9 for each of the other groups. Vect, empty vector RNAi. \**P*<0.05, \*\**P*<0.01.

Strikingly, five out of eight HSPG genes were found to be required for PFF-induced motor dysfunction (Fig. 4A). In addition to the cell surface proteoglycan syndecan/ *sdn-1*, the HSPG biosynthetic enzymes *EXT1*/ *rib-1* (exostosin glycosyl-transferase 1), *EXTL3/ rib-2* (exostosin like glycosyltransferase 3), *NDST1/ hst-1* (N-deacetylase and N-sulfotransferase 1), and *HS3ST6/ hst-3*.*2* (heparan sulfate-glucosamine 3-sulfotransferase 6) all significantly rescued motor function when knocked down by RNAi (Fig. 4A). To further test if these genes regulate susceptibility to PFF-induced aggregation of host α-synuclein, day 4 animals were imaged and α-synuclein aggregates in the muscle were quantified. Three out of these five genes when knocked down achieved a remarkable reduction in α-synuclein aggregates, both in terms of number of aggregates and total aggregate area (Fig. 4B-D). These genes, *sdn-1, rib-1*, and *hst-1*, are likely mediators of prion-like PFF seeding of host α-synuclein, whereas it is possible that the other two genes, *rib-2* and *hst-3*.*2*, which rescued motor function but did not reduce aggregate burden, may play a role in non-prion-like PFF toxicity instead.

Finally, to determine if the three HSPG genes that regulate PFF-induced toxicity and seeding of aggregation in the muscle also play a role in PFF acceleration of dopaminergic neuron degeneration, worms expressing neuronal α-synuclein were crossed to a neuronal RNAi-sensitive background and raised on HSPG gene RNAi’s starting at L4. On day 1, the worms were treated with 0.25 ng/μL PFFs, and dopaminergic neurons were assessed for neurodegeneration on day 5. All three HSPG genes that afforded protection from prion-like PFF phenotypes in worms expressing muscle α-synuclein significantly increased the number of CEP dopamine cell bodies in PFF-fed worms with neuronal α-synuclein expression (Fig. 5A-B). We did not find any effect on dopaminergic neurites (Fig. 5C), but interestingly, *hst-1* knockdown significantly increased the number of ADE cell bodies (Fig. 5D). These results suggest that *sdn-1, rib-1*, and *hst-1* play important roles in prion-like α-synuclein transmission from the digestive tract to neurons and muscle, and that the HSPG pathway may more generally protect against both prion and non-prion-like mechanisms of α-synuclein-induced injury.

**Fig. 5:**
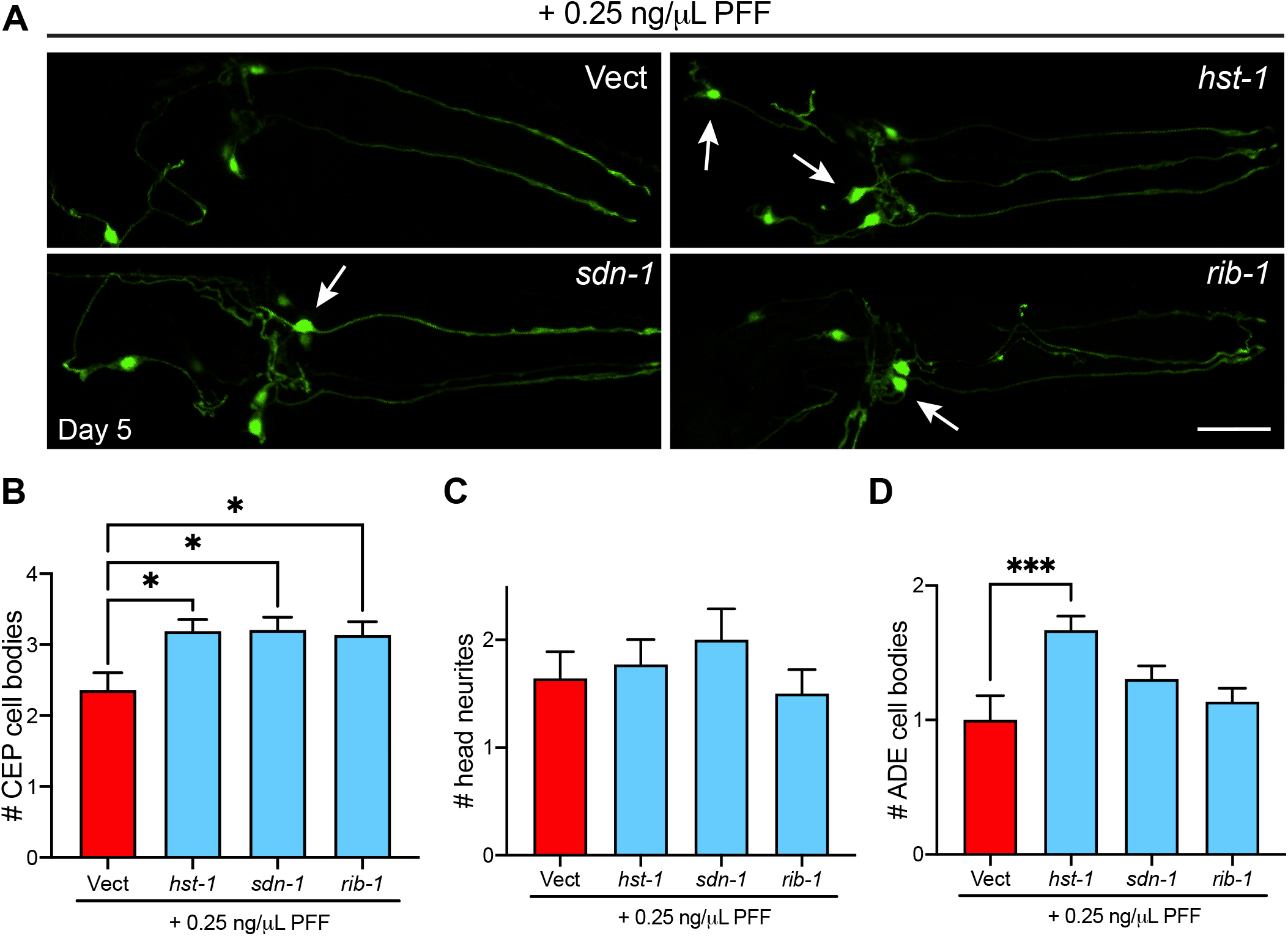
PFF-induced dopaminergic neurodegeneration is mitigated by HSPG gene knockdown. **A**, Worms expressing pan-neuronal human wild-type α-synuclein in a neuronal RNAi-sensitive background were treated with 0.25 ng/μL PFFs on day 1 of adulthood. On day 5, dopamine neurons were imaged and scored for rescue of cell bodies (arrows). Scale bar, 20 μm. **B**, CEP cell body quantification. Data are mean±s.e.m. One-Way ANOVA with Dunnett’s post-hoc. **C**, ADE cell body quantification. Data are mean±s.e.m. One-Way ANOVA with Dunnett’s post-hoc. **D**, Neurite quantification. Data are mean±s.e.m. One-Way ANOVA with Dunnett’s post-hoc. In B-D, n=14 for Vect, 22 for *rib-1*, 21 for *hst-1*, and 24 for *sdn-1*. \**P*<0.05, \*\*\**P*<0.001.

## Discussion

Here we offer new *C. elegans* models of PD that recapitulate gut-derived spreading of α-synuclein with resulting dopamine neuron degeneration and toxic seeding of host α-synuclein aggregation. In PD patients, the progressive buildup of α-synuclein pathology is consistent with prion-like α-synuclein transmission^3^, and the observation of α-synuclein positive inclusions in the enteric nervous system is suggestive of a ‘gut-to-brain’ disease progression^5^. Moreover, one of the earliest regions in the central nervous system to develop α-synuclein pathology is the dorsal motor nucleus of the vagus, which connects the brain to the periphery^4^.

Several lines of evidence from animal models support the notion that α-synuclein cell-to-cell transmission may be a critical factor in neurodegenerative disease. In rodents, intracerebral inoculation of either α-synuclein PFFs^21^, brain tissue from symptomatic A53T mutant α-synuclein transgenic mice^22^, or brain tissue from synucleinopathy patients^23^ results in widespread deposition of Lewy body-like α-synuclein inclusions often accompanied by neurodegeneration and motor deficits. Several studies have also demonstrated the spread of α-synuclein pathology from peripheral sites to the central nervous system, using intravenous^24^, intramuscular^25^, or intraperitoneal^26^ injections of recombinant α-synuclein aggregates into rodents. More recently, oral administration or gastrointestinal injection of PFFs in mice were shown to recapitulate gut-to-brain α-synuclein spread and PD-like disease^7-9^.

In *C. elegans*, a small number of studies have shown cell-to-cell transmission of α-synuclein. Using a split-GFP approach in distinct neuronal subsets, Tyson et al. showed that α-synuclein could be transferred from neuron-to-neuron^13^. A similar approach reported transfer of α-synuclein between pharyngeal muscle and neuronal cells, with disease-associated phenotypes^14^. Cross-tissue transmission of α-synuclein has also been observed from dopaminergic neurons or muscle to the hypodermis^15^. While these studies demonstrate that α-synuclein can travel between cells and even tissues in *C. elegans*, in most cases it is not known if the transmission of α-synuclein is prion-like, i.e. capable of seeding aggregation of α-synuclein in the recipient cell. Moreover, the nature of the α-synuclein species, i.e. monomers, oligomers, fibrils, or even physiological conformers, is not known due to transgenic expression of α-synuclein in both the donor and recipient cells.

In contrast, our study uses biochemically well-defined aggregated species of α-synuclein as the initial seed, and demonstrates the induction of aggregation of host α-synuclein in a prion-like manner. In a recent study in which α-synuclein oligomers were fed to *C. elegans* and were able to cross the intestinal barrier and cause toxicity^27^, the worms lacked host α-synuclein and thus these are not models of prion-like α-synuclein activity but rather provide insights into non-prion-like mechanisms of α-synuclein-initiated disease. In our models, we found that later time-points following PFF feeding, or exposure to high PFF concentrations, engaged non-prion-like mechanisms which might be similar to those induced by exogenous oligomers. In addition, we observed weak toxicity of monomeric α-synuclein, which has been reported in mice as well^28^, and remains to be fully understood.

PFF feeding in *C. elegans* provides a new approach for the modeling of PD phenotypes and disease progression. The vast array of tools available for genetic manipulation and the feasibility of large-scale investigations using *C. elegans* makes it an advantageous model system for the discovery of new disease mechanisms and potential therapeutic targets. We have explored the role of the HSPG pathway in potentially mediating PFF neuronal and muscle cell injury in *C. elegans*, and identified syndecan/ *sdn-1* as well as particular enzymes involved in polymerizing and modifying chains of sugar residues during HSPG synthesis as being important for PFF disease processes. *sdn-1* has known roles in nervous system development and axon guidance^29^, and may serve as a potential PFF receptor in our models. In addition, the specific involvement of some HSPG synthesis enzymes and not others in PFF disease phenotypes suggests that specific modification patterns may be required for PFF cellular uptake. Together, these findings open up new avenues of investigation in a versatile system for further discovery.

## Materials and Methods

### *C. elegans* strains

Worms were maintained at 20°C on standard nematode growth medium (NGM) plates or high growth medium (HG) plates, seeded with OP50 *E. coli* or HT115 RNAi *E. coli*, as indicated. The following strains were used in this study: wild-type N2 Bristol strain, NL5901 pkIs2386 (*unc-54p::hWTα-synuclein::yfp*), LC108 uIs69 [pCFJ90 (*myo-2p::mCherry + unc-119p::sid-1*)], UM0011 (*dat-1p::gfp; aex-3p::hWTα-synuclein*), AM134 rmIs126 (*unc-54p::yfp*), BY250 (*dat-1p::gfp*), and MOR002 [*dat-1p::gfp; aex-3p::hWTα-synuclein*]; uIs69 [pCFJ90 (*myo-2p::mCherry + unc-119p::sid-1*)] obtained by crossing UM0011 with LC108.

### α-Synuclein PFF preparation and treatment

Recombinant human wild-type α-synuclein (Proteos) was incubated at 1 mg/ml at 37 °C and shaking at 1,400 rpm for up to 10 d. Fibrillization was assayed by Thioflavin T (Sigma) added to a final concentration of 25 µM. Fluorescence emission was measured at 482 nm during excitation at 450 nm. Sedimentation analysis was performed by centrifugation at 16,000 g for 10 min at 4°C. The supernatants and pellets were boiled in SDS sample buffer at 95°C for 10 min, and run on SDS-PAGE gels (Invitrogen). The gels were then stained with Coomassie Blue R-250. When α-synuclein fibrillization had reached the plateau phase, and insoluble α-synuclein was present, PFFs were aliquoted and stored at -80°C until use. On the day of *C. elegans* treatment, PFFs were thawed at room temp, diluted to 0.1 mg/mL in PBS, and sonicated. Freshly sonicated PFFs were then mixed with OP50 at the indicated concentrations in a total volume of 300 μL M9, dispensed onto unseeded NGM plates and allowed to dry for 1-2 hours prior to transferring to the plates day 1 adult worms that had been synchronized by bleaching. Monomer α-synuclein was prepared exactly in parallel, except with thawing on ice and without sonication. Buffer treatments used M9 in the absence of α-synuclein. The day after treatment, worms were transferred to new NGM plates that had been seeded with OP50 without α-synuclein. Thereafter, worms were transferred to new OP50-seeded NGM plates without α-synuclein every 2 days for the duration of the experiment.

### RNAi treatments

Worms were synchronized by bleaching and plating onto HG plates seeded with OP50. At the L4 stage, worms were transferred to RNAi-seeded NGM plates containing carbenicillin and IPTG that were pre-induced with 0.1 M IPTG. The next day, freshly sonicated PFFs were mixed with RNAi bacteria at the indicated concentrations in a total volume of 300 μL M9, dispensed onto unseeded RNAi plates, induced with 0.1 M IPTG, and allowed to dry for 1-2 hours prior to transferring the day 1 worms. Buffer treatments used M9 in the absence of α-synuclein. The day after treatment, worms were transferred to new RNAi plates that had been seeded without α-synuclein and were pre-induced with 0.1 M IPTG. Thereafter, worms were transferred to new RNAi plates without α-synuclein every 2 days for the duration of the experiment. Control vector RNAi refers to empty vector pL4440 in HT115 *E. coli*.

### Neurodegeneration imaging

Worms expressing GFP in dopamine neurons (strains UM0011, MOR002, and BY250) were mounted on 2% agarose pads with M9 and sodium azide, and imaged on a Nikon A1R MP+ multiphoton/confocal microscope at 60x magnification. To count cell bodies, Z stacks were analyzed using Fiji software. Maximum intensity projections were generated and a region of interest was drawn around each cell body. A numerical threshold for area (μm^2^) was set to distinguish cell body size that would be considered present/non-degenerated. Once a threshold was set, it was used across all treatment groups in the experiment. To avoid errors in measuring neurons that appeared colocalized in the maximum intensity projection, the original Z stack was used to identify and measure each neuron separately. All visible ADEs and CEPs were measured. Neurites projecting from CEP cell bodies were scored using 3D rendering in NIS Elements software. Neurites were counted as present if they did not have degenerated morphologies such as blebbing or fragmentation.

### α-Synuclein aggregation imaging

Worms expressing α-synuclein-YFP in muscle cells (strain NL5901) or YFP in muscle (strain AM134) were mounted on 2% agarose pads with M9 and sodium azide, and imaged on a Nikon A1R MP+ multiphoton/confocal microscope at 60x magnification. Z stacks were analyzed using Fiji software. Maximum intensity projections were generated and an intensity threshold was set in order to distinguish aggregates from background. Once a threshold was set, it was used across all treatment groups in the experiment.

### Motor assays

At indicated ages, worms (strains NL5901 or N2) were individually picked onto unseeded NGM plates. Worms were allowed to recover for 1 min, and then the number of body bends was manually counted for 30 sec.

### Statistical analysis

Statistical analyses were performed using GraphPad Prism 9. An unpaired two-tailed Student’s *t*-test was used for all comparisons between two groups. For comparisons between multiple groups, One-Way ANOVA or Two-Way ANOVA (for repeated measures or two-variable analyses, i.e. motor assays with multiple treatment groups and multiple strains) was performed with post-hoc testing as indicated.

## Acknowledgments

We thank the *C. elegans* Genetics Center for strains (P40 OD010440), and the Department of Neuroscience and Regenerative Medicine 2-Photon Microscopy Facility. Strains N2 and NL5901 were generously provided by C. Murphy (Princeton University). Strain UM0011 was generously provided by G. Wong (University of Macau) and C. Wang (University of Hong Kong). Strain BY250 was generously provided by R. Blakely (Florida Atlantic University). Strain AM134 was generously provided by R. Morimoto (Northwestern University). This work was supported by the start-up fund from the Medical College of Georgia at Augusta University.

## Author Contributions

D.E.M., M.C., and J.V. conceived and designed the experiments. M.C., J.V., A.E., S.W., S.Y. and D.E.M. performed the experiments and analyzed the data. M.C. and D.E.M. wrote the paper.

---

**Fig S1:**
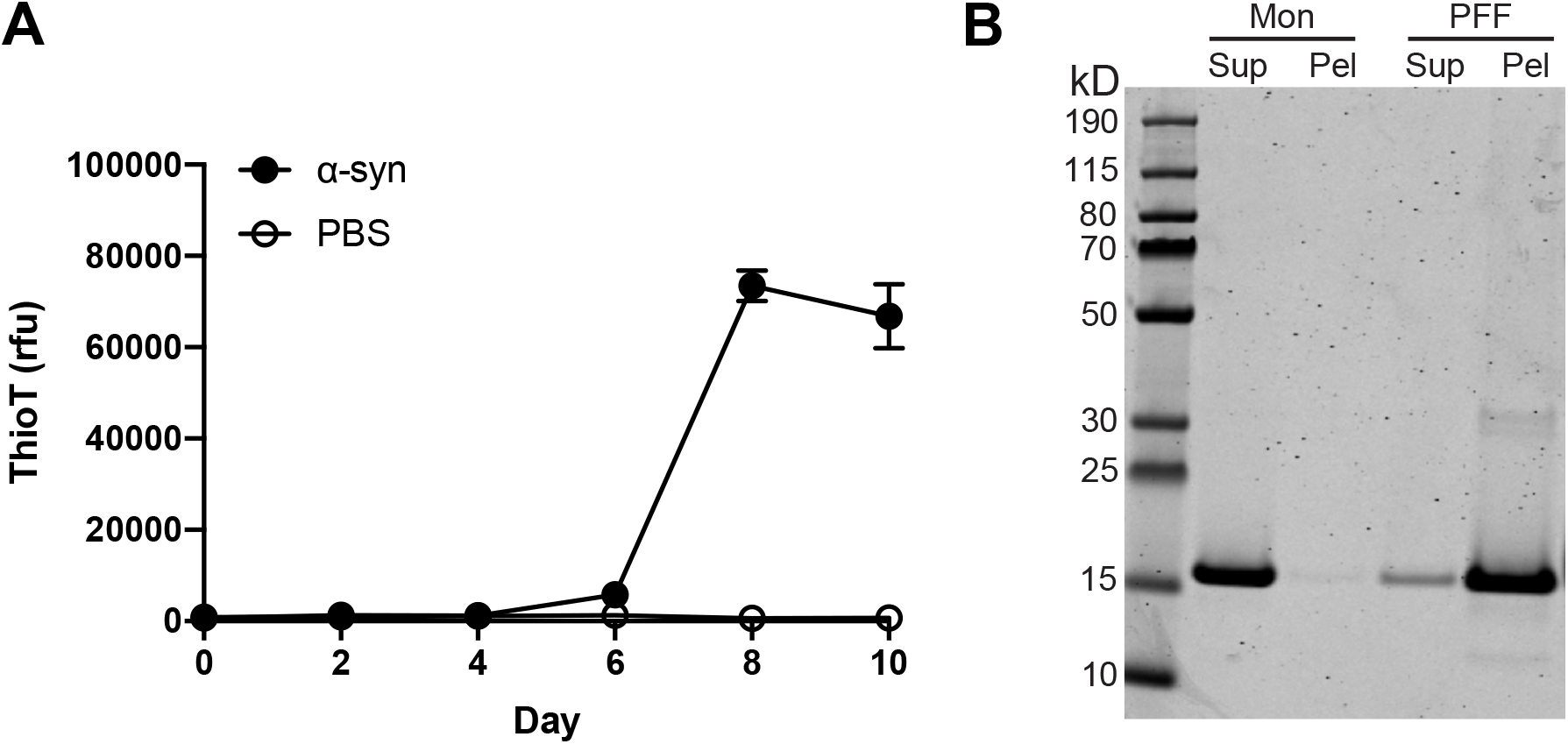
PFF characterization. **A**, Thioflavin T curve showing recombinant α-synuclein protein fibrillization (1 mg/mL). n=3 wells per group. rfu, relative fluorescence units. **B**, SDS-PAGE showing α-synuclein in the supernatant (Sup) and pellet (Pel) for monomer and PFF samples.

